# Isolation of proteins on chromatin (iPOC) reveals signaling pathway-dependent alterations in the DNA-bound proteome

**DOI:** 10.1101/2024.04.10.588873

**Authors:** Huiyu Wang, Jeroen Krijgsveld, Gianluca Sigismondo

## Abstract

Signaling pathways often convergence on transcription factors (TFs) and other DNA-binding proteins (DBPs) that regulate chromatin structure and gene expression, thereby governing a broad range of essential cellular functions. However, the repertoire of DBPs is incompletely understood even for the best-characterized pathways. Here, we optimized a strategy for the isolation of Proteins on Chromatin (iPOC) exploiting tagged nucleoside analogues to label the DNA and capture associated proteins, thus enabling the comprehensive, sensitive, and unbiased characterization of the DNA-bound proteome. We then applied iPOC to investigate chromatome changes upon perturbation of the cancer-relevant PI3K/AKT/mTOR pathway. Our results show distinct dynamics of the DNA-bound proteome upon selective inhibition of PI3K, AKT, or mTOR, and we provide evidence how this signaling cascade regulates the DNA-bound status of SUZ12, thereby modulating H3K27me3 levels. Collectively, iPOC is a powerful approach to study the composition of the DNA-bound proteome operating downstream of signaling cascades, thereby both expanding our knowledge of the mechanism of action of the pathway, and unveiling novel chromatin modulators that can potentially be targeted pharmacologically.

## 1. Introduction

In eukaryotic cells, chromatin is a tightly packed and organized complex mainly composed of DNA and histone proteins [1]. In addition, many non-histone proteins dynamically associate with chromatin to fine-tune gene expression programs in response to intracellular and extracellular signals, thus regulating a wide range of biological processes [2]. Moreover, a number of protein machineries associate with chromatin to repair DNA damage inflicted by transcriptional stress or environmental cues, thereby safeguarding genomic integrity [3]. To fulfill all of these tasks, chromatin-associated proteins span a wide range of functionalities including transcription factors, epigenetic writers, readers and erasers (both of histones and DNA), and proteins that detect and repair various types of DNA damage [4]. Of note, these factors often need to act in concert in a highly ordered fashion to maintain cellular homeostasis. Due to the key role of chromatin-binding proteins, alterations in expression and chromatin association of chromatin regulators are observed in many human diseases [5]. For instance, neurodegenerative and other neurological disorders are generally characterized by histone hypo-acetylation due to reduced histone acetyltransferase (HAT) activity or increased histone de-acetylase (HDAC) function [6]. In addition, about 50% of the commonly identified cancer driver genes encode for chromatin-bound proteins, strongly highlighting the impact of chromatin composition dysregulation in cancer development [7]. Elective examples include the SWI/SNF components, a subfamily of ATP-dependent chromatin remodeling complexes mutated in 20% of all human cancers [8], the protein TP53 that corresponds to the most commonly mutated gene across the Pan-Cancer collection [9], and subunits of the Polycomb repressive complex 2 (PRC2), strongly linked to cancer progression. In view of their disease-relevant role, chromatin proteins are desirable targets of epigenetic drugs[10, 11], and therefore the comprehensive identification of DNA-binding proteins could contribute to identify new regulators of cellular functions, that could serve as potential novel drug targets.

The composition of the DNA-bound proteome (i.e. chromatome) is often orchestrated by signaling cascades that function by connecting distinct upstream stimuli (e.g. growth factors) with an alteration in chromatin composition (e.g. activation of a TF) to induce the expression of a specific repertoire of genes, thus ultimately executing the cellular response (e.g. proliferation) of the physiological or pathological insult. Among the pathways that are important for maintaining cellular homeostasis, the PI3K/AKT/mTOR signaling responds to nutrients, hormones and growth factors, by promoting the phosphorylation of a number of key players of fundamental biological processes such as FOXO, CHK1, p21, and GSK3, thus ultimately regulating cell growth, motility, survival, and metabolism. Deregulation of the PI3K/AKT/mTOR signaling cascade leads to increased survival and cell cycle progression, and is therefore observed in nearly all types of neoplasm, ranging from colorectal carcinoma and breast cancer, to non-small cell lung cancer, and ovarian cancer [12]. To curb these deleterious processes, various inhibitors targeting selective components of these signaling pathway have been developed as cancer therapy [13]. Among the inhibitors of the PI3K/AKT/mTOR axis, Taselisib, MK-2206, and Torin1 specifically act on PI3K, AKT and mTOR, respectively [14–16]. Despite the initial efficacy of these drugs, target cells often develop resistance by either acquiring hyperactivating mutations of factors downstream of the drug target, or by compensatory effects of other signaling cascades (e.g. MEK/ERK pathway) [17]. The comprehensive understanding of therapy failure or development of resistance is hampered by the limited knowledge of downstream effectors of this pathway acting on DNA. Moreover, very little is known about whether PI3K, AKT, and mTOR trigger a similar regulation of chromatin. For this reason, while phospho-proteomic studies focused on merely dissecting a signaling cascade fall short in identifying the ultimate readout of pathway activation, the study of the modulation of the DNA-bound proteome might lead to the potential identification of additional druggable targets that could be used in combination therapy.

The study of the DNA-bound proteome requires the purification of chromatin to high homogeneity; however, this is challenging due to its water-insoluble and highly charged nature which resists to mechanical, enzymatic, and ion-based extraction. In addition, separation of chromatin from other cellular organelles is difficult, often resulting in contamination with cytosolic proteins [3, 18]. Yet, several biochemical strategies have been recently developed that effectively combine different means of chromatin purification strategies with mass spectrometry-based protein identification [19]. In this regard, crosslinking strategies to stabilize protein-DNA interactions allow for stringent washing conditions and resulting in higher chromatin purity, e.g. as used in Chromatin Enrichment for Proteomics (ChEP) [20]. To further boost the sensitivity of DNA-bound proteome identification, a variety of strategies have been developed that take advantage of nucleoside analogue integration and click chemistry-based enrichment of DNA. Successful examples are the isolation of proteins on nascent DNA (iPOND) and DNA-mediated chromatin pulldown (Dm-ChP) [19, 21, 22], where alkyne-containing thymidine analogs (e.g. 5-ethynyl-2′-deoxyuridine, EdU) are incorporated into replicating DNA. Azide biotin is then covalently linked to the DNA by click chemistry reaction, followed by streptavidin-based enrichment of the DNA-bound proteome of replicating DNA. Similarly, more extensive integration of EdU into DNA can be exploited for the identification of protein on total DNA (iPOTD) [23]. Inspired by these technologies, we recently refined this concept and combined it with our recently developed protease-resistant streptavidin beads for the ultra-sensitive isolation of protein on chromatin (iPOC), and we successfully applied it to characterize the dynamics of the DNA-bound proteome during DNA double-strand break (DSB) repair [24].

Here we optimized the iPOC protocol by defining the optimal concentration of EdU and protease-resistant streptavidin beads, and by determining best decrosslinking, digestion and desalting conditions. After thorough assessment of the specificity and sensitivity of the approach, we apply iPOC to the dissection of the DNA-bound proteome upon selective inhibition of PI3K, AKT, or mTOR. Our results demonstrate the benefit of iPOC by shedding light on previously unknown common as well as unique effects of PI3K-, AKT-, and mTOR-inhibition on chromatin composition.

## 2. Method

### 2.1 Experimental design and statistical rationale

For the optimization of iPOC, MCF7 cells were treated with 5-ethynyl-2’-deoxyuridine (EdU) for 18h with a final concentration of 0.1, 0.2, 1, 10, 20, or 50 μM each in three replicates. For the optimization of sample preparation, DNA-bound complexes were de-crosslinked for 5min or 20min at 95℃ on protease-resistant streptavidin (prS) magnetic beads [25]. After reduction and alkylation, both the supernatant and the bead were digested overnight with trypsin, followed by peptide clean-up by either SP3 (single-pot, solid phase-enhanced sample preparation) [26] or C18-stage tip protocol [27]. For the competition experiment, 100%, 75%, 25%, 0% biotin-azide was competed with 0%, 25%, 75%, 100% cy5-azide in EdU-labeled cells. Each condition was performed in two biological replicates. For the experiments with drug treatments, we combined the optimized iPOC procedure with triple SILAC labeling and mass spectrometry analysis to quantify dynamic changes in the DNA-bound protein composition induced by inhibition of PI3K, AKT, or mTOR signaling pathways in three biological replicates. In particular, SILAC-intermediate-labeled MCF7 cells were only exposed to EdU as internal control, while heavy-labeled cells were treated with both EdU and either PI3K, AKT, or mTOR inhibitor. Light-SILAC cells were treated with DMSO and served as negative control.

### 2.2 Cell culture and SILAC labeling

MCF7 cell lines were cultured in DMEM medium with 10% fetal bovine serum and 1% penicillin-streptomycin. Cells were incubated at 37 °C with CO_2_ level of 5% and split at 90% confluency. For SILAC labeling, cells were metabolically labeled in SILAC-compatible DMEM medium containing light (Arg0, Lys0), intermediate (Arg6, Lys4) or heavy (Arg10, Lys8) amino acids. Light-labeled samples were collected in untreated conditions, intermediate-labeled samples were treated with EdU for 18h, while heavy labeled cells were treated with EdU for 18h and exposed for 6h to PI3K, AKT, mTOR inhibitor Taselisib (IC50=7.3uM), MK2206 (IC50=6.4uM), Torin 1 (IC50=3uM), respectively.

### 2.3 Isolation of protein on chromatin (iPOC)

For each experiment, 4×10^7^cells were treated with 5-ethynyl-2’-deoxyuridine (EdU) for 18 h at a concentration of 0.1, 0.2, 1, 10, 20, or 50 μM. Cells were then crosslinked with 1% formaldehyde in PBS for 10 min and quenched for 5 min with glycine at a final concentration of 125mM. After two washes with 1x PBS, cells were resuspended in 4 ml of permeabilization buffer (0.25% Triton X-100) and incubated at RT for 30 min. Cells were then spun down for 5 min at 900g and washed once with cold 0.5% BSA in 1x PBS, and once in 1x PBS. Permeabilized cells were then resuspend in click reaction buffer (10 μM biotin azide (Jena Bioscience CLK-047), 10 mM sodium ascorbate, 2 mM CuSO_4_ in PBS 1%) at RT, in the dark for 2h. Cells were then spun down for 5 min at 900g, washed once with cold 0.5% BSA in PBS, and once with 1x PBS. Cells were then lysed in 300 μl of lysis buffer (1% SDS, 50 mM pH 7.5 Tris-HCl, protease inhibitor) and sonicated with the Pico Bioruptor (Diagenode) for 12 cycles (30s ON/ 30s OFF). Samples were centrifuged for 10 min at 16,000g and protein concentration was measured by BCA assay (Thermo Scientific, 23225). For SILAC labelling, an equal amount of supernatants were taken before adding 100 μl of prS beads [25] pre-conditioned with lysis buffer. Samples were rotated at RT for 3h and beads with chromatin complex were collected on the magnetic rack. After washing once with lysis buffer and once 1M NaCl, streptavidin-enriched DNA-bound complexes were resuspended in 20μl of 50mM ammonium bicarbonate (AmBic) decrosslinked at 95℃, reduced with 10mM DTT for 30min at 45℃, and alkylated with 20mM iodoacetamide for 30min in dark. Proteins were finally digested on beads with 500ng of trypsin (Promega, V5280) for 18 h. Peptides were cleaned up using either SP3 [26] or C18-stage tip protocol [27], reconstituted in 0.1% trifluoroacetic acid (TFA), and stored at - 20℃ till mass spectrometry analysis.

### 2.4 TMT labeling

The peptides from competition experiment were labeled with the TMT10-plex reagent (126, 127 N, 127 C, 128 N, 128 C, 129 N, 129 C, 130 N, 130 C, 131) performed according to manufactureŕs instructions (Thermo Scientific, 90110). Briefly, peptides from 10 samples were suspended in 20μl of 50mM TEAB buffer (pH 8.5). Then, 0.8mg of TMT reagent in 41 μl of acetonitrile was added to each sample, mixed thoroughly, and incubated at 25°C for 1h. After that, 8μl of 5% Hydroxylamine solution was added to each sample to stop the reaction. Finally, the labeled peptides were pooled and purified by the OASIS C18 Cartridge (Waters, 186008052). Cleaned peptides were eluted twice by ethanol 70%, dried in SpeedVac and stored at −80 °C until high-pH fractionation.

### 2.5 High-pH peptide fractionation

Ammonium formate with final concentration 20mM was added to each sample before fractionation *via* high-pH reverse-phase chromatography. Peptides were fractionated on an Agilent 1290 Infinity HPLC system (Agilent) with a Gemini C18 column (3 μm, 110Å, 100 × 1.0 mm, Phenomenex) using a linear 60 min gradient from 0% to 35% (v/v) acetonitrile in 20 mM ammonium formate (pH 10) at a flow rate of 0.1 ml/min. Elution of peptides was detected with a variable-wavelength UV detector set to 254 nm. A total of 40 fractions were collected and subsequently concatenated into 8 final fractions per sample.

### 2.6 Mass spectrometry data acquisition

Peptides were loaded on a trap column (PepMap100 C18Nano-Trap 100 μm × 20 mm) and separated over a 25cm analytical column (Waters nanoEase BEH, 75 m × 250mm, C18, 1.7μm, 130Å) using a Thermo Easy nLC 1200 HPLC system (Thermo Fisher Scientific). Solvent A was water with 0.1% formic acid, and solvent B was 80% acetonitrile, 0.1% formic acid. During the elution step, the percentage of solvent B increased in a linear fashion from 3 to 8% in 4 min, then increased to 10% in 2 min, to 32% in 68 min, to 50% in 12 min and finally to 100% in a further 1min and went down to 3% for the last 11min. Peptides were analyzed on a Tri-Hybrid Orbitrap Fusion mass spectrometer (Thermo Fisher Scientific) operated in positive (+2.5 kV) data dependent acquisition mode with HCD fragmentation. The MS1 and MS2 scans were acquired in the Orbitrap and ion trap, respectively, with a total cycle time of 3s. MS1 detection occurred at 120 000 resolution, AGC target 1E6, maximal injection time 50ms and a scan range of 375-1500 m/z. Peptides with charge states 2-4 were selected for fragmentation with an exclusion duration of 40s. MS2 occurred with NCE 33%, detection in topN mode and scan rate was set to Rapid. AGC target was 1E4 and maximal injection time allowed of 50 ms. Data were recorded in centroid mode. For TMT experiment, TMT samples were quantified at MS2 level.

### 2.7 Proteomic data processing

Raw data were analyzed by MaxQuant software (v2.4.0.0) [28]by searching against the UniProtKB human database (release 2016_11_28, 20244 entries). The database search followed an enzymatic cleavage rule of Trypsin/P, allowed maximal two missed cleavage sites, and a mass tolerance of 20 ppm for fragment ions. Carbamidomethylation of cysteines was defined as fixed modification, while N-terminal acetylation, methionine oxidation were defined as variable modifications. Maximum FDR was set at 1% for both peptides and proteins level. Perseus software was used for data visualization [29]. After canonical filtering (reverse, potential contaminants and proteins only identified by site), only proteins with at least 2 unique peptide in all the replicates were considered as identified while only proteins with LFQ or SILAC ratio in all the replicates were defined as quantified. Pathway enrichment analysis was performed using the Metascape web software [30].

For analysis of TMT-labeled samples, TMT10plex with reporter at MS2 level was specified in MaxQuant search. Total intensity of TMT intensity in each sample was normalised before data processing to account for potential mixing error.

### 2.8 Western blot

For iPOC, cells were collected before and after the click chemistry reaction as pre-labeling and post-labeling. The samples before and after streptavidin enrichment were collected as input and flow-through, respectively, to evaluate efficiency of streptavidin enrichment. For inhibitor experiments, MCF cells were treated with the specific inhibitor for 6h, 12h, or 24h, and harvested in RIPA buffer (10 mM Tris–HCl pH 8.0, 140 mM NaCl, 1 mM EDTA, 0.1% SDS, 1% Triton 100) with Phosphatase protease inhibitors. The cell lysates were then sonicated (3 pluse, 10s, 10% amplitude) and centrifuge at 12000 g for 10 min. Then, the supernatant was transferred to a new tube, and protein concentration was measured by BCA assay (Thermo Scientific, 23225).

Twenty μg of protein extracts were separated on SDS–PAGE followed by electrotransfer to a PVDF membrane. Membranes were blocked in 5% non-fat milk for 1h and then incubated overnight at 4°C with primary antibodies: PI3K p85 alpha (#ab191606, 1:1,000, abcam), Phospho-PI3K p85 alpha (#ab182651, 1:1,000, abcam), AKT (# ab179463, 1:1,000, abcam), Phospho-AKT (ab192623, 1:1,000, abcam), Phospho-mTOR (Ser2448) (#5526, 1:1,000, Cell Signaling Technology), mTOR (#2983, 1:1,000, Cell Signaling Technology), Phospho-FoxO1 (#9461, 1:1,000, Cell Signaling Technology), FoxO1 (#2880, 1:1,000, Cell Signaling Technology), β-actin (#4970, 1:5,000, Cell Signaling Technology), Histone H2A (ab18255, 1:1,000, Abcam), SUZ12(#3737, 1:1,000, Cell Signaling Technology), H3K27me3(#9733, 1:1,000, Cell Signaling Technology), HRP-conjugated streptavidin (#3999, 1:2000, Cell Signaling Technology). The membranes were then washed with washing buffer (Tween 0.1% in 1x PBS) and incubated with HRP-conjugated anti-rabbit IgG secondary antibody for 1h at room temperature. Protein abundance were detected by the ECL™ Western Blotting Detection Reagents (GE Healthcare) and visualized on a Chemidoc system (BioRad). Westernblot quantification analysis was performed by ImageJ software [31].

## 3 Results

### 3.1 Optimization of EdU concentration in iPOC procedure

We recently introduced the strategy for isolation of protein on chromatin (iPOC) where DNA is labeled with the thymidine analog 5-ethynyl-2′-deoxyuridine (EdU), followed by click chemistry-based DNA tagging, and enrichment of stabilized DNA-bound proteins (DBPs) on protease-resistant streptavidin beads [24]. We effectively employed iPOC to capture and identify the differential association of DBPs during the DNA damage response [24]. Here we aimed to further optimize the iPOC protocol by systematically evaluating the experimental conditions with regard to EdU labeling, protein decrosslinking, digestion and peptide clean-up.

We first focused on finding the minimal concentration of EdU that would allow for whole DNA labeling, while ensuring the highest protein recovery. We therefore performed iPOC in MCF7 cells exposed to increasing concentrations of EdU for 18h, and evaluated the labeling efficiency, as well as the number of proteins identified *via* MS upon streptavidin enrichment. EdU-labeled cells were then treated with 1% formaldehyde to stabilize protein-DNA interactions, followed by cell permeabilization and copper-catalyzed azide-alkyne cycloaddition (CuAAC) between biotin-azide and EdU-labeled DNA. Upon cell lysis and chromatin preparation, biotin-labeled DNA– protein complexes were purified using protease-resistant streptavidin (prS) beads [25], and subjected to tryptic digestion for the MS-based proteomic investigation of the DNA-bound chromatin protein composition (Fig. 1A).

**Figure 1.**
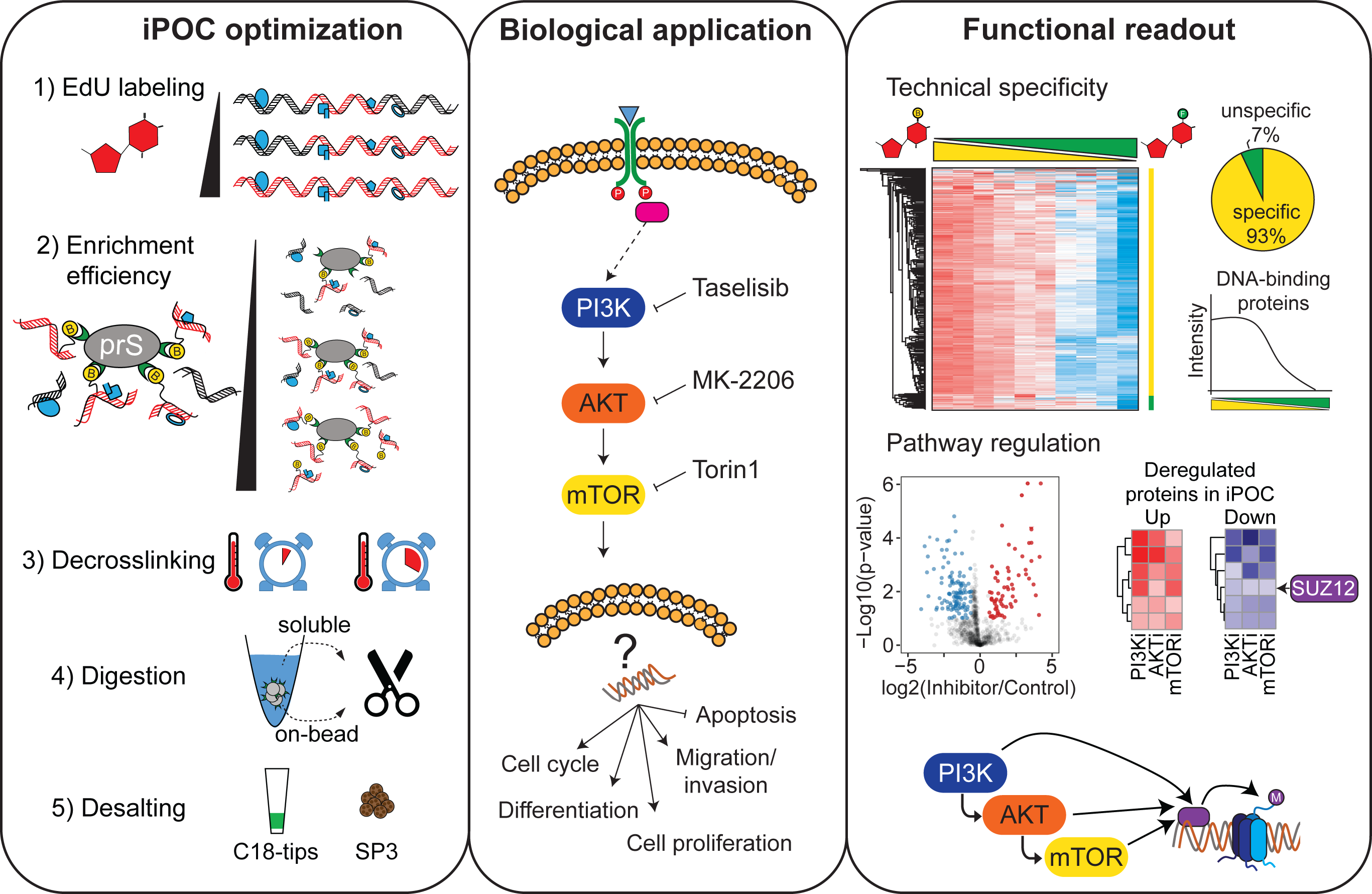
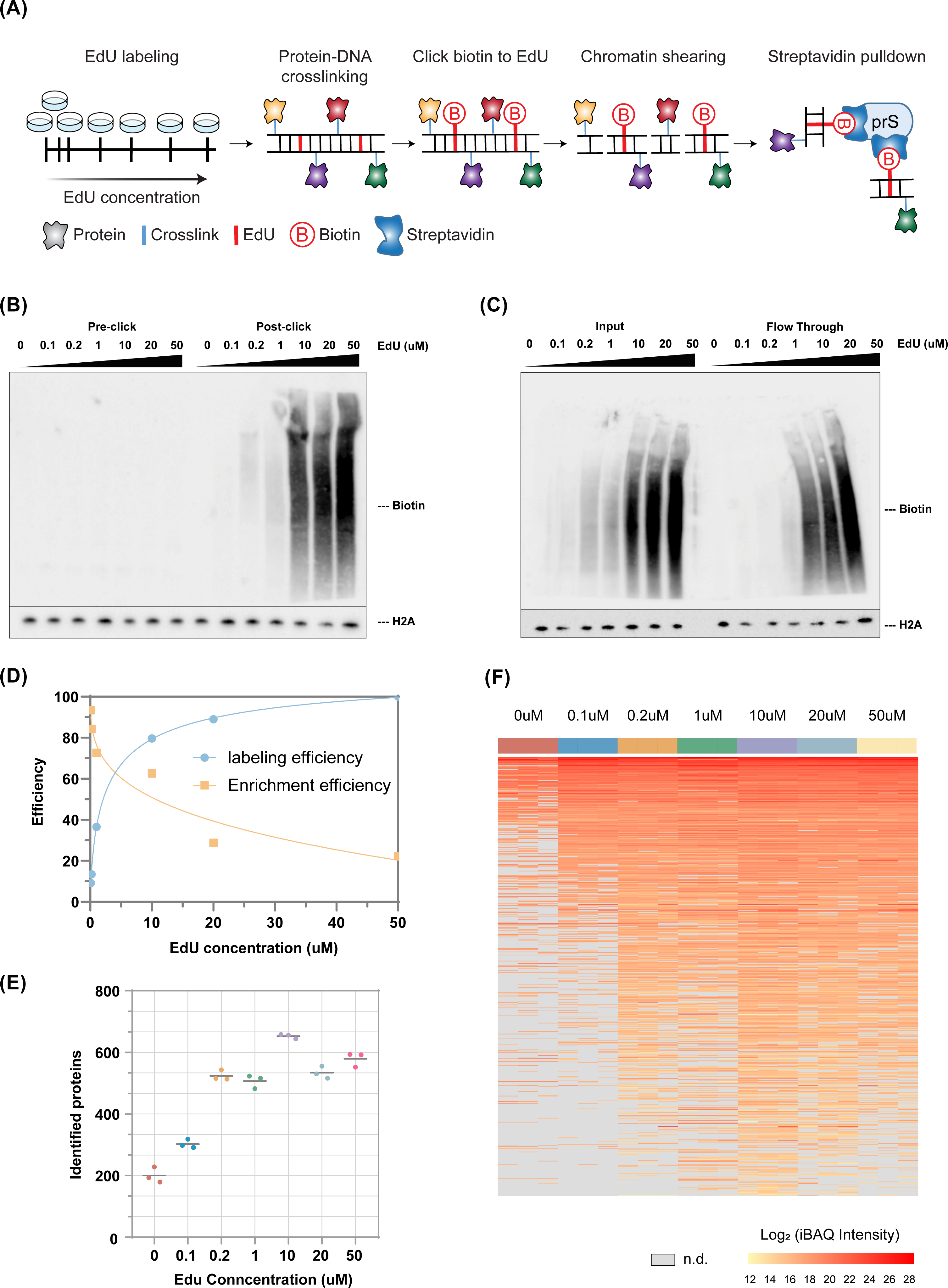
Optimization of EdU concentration in iPOC procedure. (A) Schematic of iPOC experimental design. (B) Western blot showing biotin (top) and histone H2A (as normalizer (bottom)) before (Pre-click) or after (Post-click) click reaction with biotin azide at increasing EdU concentrations. (C) Western blot showing biotin (top) and histone H2A (as normalizer (bottom)) in samples before (Input) or after (Flow through) protease-resistant streptavidin (prS)-mediated enrichment of DNA-bound proteins. (D) Line plot representing the quantification of labeling and enrichment efficiencies from B and C, as well as respective fitting curves. (E) Number of proteins identified *via* mass spectrometry in iPOC experiments carried out with cells exposed at different concentrations of EdU (n=3). (F) Heatmap representation showing the iBAQ intensity of proteins identified in iPOC at different EdU concentrations. Protein intensity is shown from yellow to red. n.d.: proteins not detected, show in grey.

To evaluate the biotin labeling efficiency, we determined the amount of biotin in the chromatin preparations from cells exposed to increasing concentrations of EdU (Post-click), relative to the respective untreated samples (Pre-click) (Fig. 1B). Western blot analysis suggests that by using equal amounts of biotin at increasing EdU concentration, the biotin labeling efficiency follows a logarithmic increase, and reaches a plateau around 10μM of EdU (Fig. 1B, Fig. 1D). In parallel, we assessed the streptavidin enrichment efficiency by comparing the signal of biotin in respective conditions before (Input) or after (Flow-through) the streptavidin enrichment (Fig. 1C). Our results suggest that, keeping the amount of prS beads constant, the efficiency of enrichment decreases exponentially at increasing concentrations of EdU (Fig. 1C, Fig. 1 D). Interpolation of these results indicates 5μM of EdU as the optimal condition in terms of both biotin labeling and streptavidin enrichment efficiency (Fig. 1D).

To investigate this in more detail, we performed a single-shot LCMS analysis of a limited amount of each iPOC sample corresponding to ∼5E6 cells, and identified almost 600 DNA-bound proteins in the three replicates at EdU concentrations ranging from 0.2 to 50 μM (Fig. 1E, Fig. 1F). Since the number of proteins saturates above 1 μM EdU (Fig. 1E), we conclude that a EdU concentration between 1 to 10 μM suffices for best results, aligning with our previous result (Fig. 1D). To investigate this further, we compared the number of proteins identified in iPOC performed in cells exposed to 1, 5, or 10μM EdU. Our results showed that the number of proteins identified at 5μM EdU was slightly higher to that of 10μM condition, while identifying more unique proteins (Fig. S1A, Fig. S1B). Accordingly, while several DNA-binding proteins were identified in all EdU-treated samples, such as zinc figure-containing proteins (ZC3H14, ZMYM2, ZNF638), known transcription factors (e.g. GATAD2A), and the transcription activator BRG1 (SMARCA4), additional chromatin regulators like lysine methyltransferase 2A (KMT2A), nucleolar transcription factor 1 (UBTF), and DNA replication licensing factors (MCM5, MCM6), were exclusively identified in cells exposed to 5μM of EdU. Due to the highest number of identified proteins and the overall enrichment of DNA-binding proteins, we therefore established 5μM of EdU as optimal working condition for the subsequent experiments. Of note, this is 4-fold lower than in our original version of the protocol[24].

### 3.2 Optimized decrosslinking, digestion and desalting strategy for efficient iPOC sample processing

To further increase the sensitivity of the approach we optimized three additional key steps in sample preparation for iPOC: protein decrosslinking/elution, digestion, and desalting (Fig. 2A). In particular, capturing on streptavidin beads allows for stringent washes, but requires harsh elution conditions to release the DNA-protein complexes. In addition, this step is also needed for the simultaneously formaldehyde de-crosslink to achieve high proteome coverage in LCMS. Since harsh elution inevitably leads to leakage of streptavidin from the beads, thereby producing a massive sample contamination, in iPOC we have largely mitigated this effect by taking advantage of our protease-resistant streptavidin (prS) beads, which prevents proteolytic cleavage of streptavidin and reduces its contamination over 100 fold, thus boosting the identification of low abundance proteins [25]. To further investigate optimal elution conditions in iPOC we compared the number of identified proteins and the abundance of streptavidin peptides upon heating the prS beads at 95℃ for either 20min or 5min to de-crosslink the chromatin and elute the DNA-binding proteins. Our results showed that increased elution time does not improve the number of identified proteins significantly, but increases the abundance of streptavidin peptides from 5% to roughly 10% of the total ion chromatogram (Fig.2B, Fig. S2A). Hence, de-crosslinking for 5 min suffices to balance high proteome coverage with low streptavidin contamination. To maximize the yield of DNA-binding proteins, we next investigated whether, under the selected decrosslinking conditions, DNA-bound proteins were effectively eluted or still remained bound to the prS beads. To test this, we separately digested the proteins eluted upon the de-crosslinking step and proteins remaining on prS beads, and evaluated their abundance *via* mass spectrometry. Irrespective of the elution conditions, our results showed that more proteins were identified by performing on-prS bead digestion (Fig.2C top, Fig.S2C). In order to compare the protein recovery upon either on-bead or in-solution digestion, we used the same amount of trypsin, and calculated the contribution of trypsin to the overall intensity by dividing the extracted ion current of the 3 most intense trypsin peptide by the base peak. Our results show that trypsin accounted for more than 60% of the total ion chromatogram upon de-crosslinking, while this percentage drastically decreased to 25% when performing on-prS beads digestion (Fig.2C bottom, Fig.2E). These results are therefore consistent with the higher number of proteins identified when digesting on-prS beads. Lastly, we compared peptide desalting with either SP3 [26] or C18 stage-tip [27]procedure, and evaluated the number of identified proteins in MS analysis as well as the overall abundance of selected DNA-bound proteins. Our result shows that there is not a significant difference in the number of identified proteins, making both clean-up methods equally efficient (Fig. 2D top). Nevertheless, we observed a slightly higher abundance of DNA-related proteins upon SP3 desalting (Fig.2D bottom, Fig. S2B). Collectively, after comparing different sample processing procedures, our results show that the best-performing protocol for the identification of DNA-binding proteins in iPOC consists of de-crosslinking at 95℃ for 5mins, tryptic digestion on the prS beads, followed by SP3 peptide desalting. These conditions were employed in the next experiments.

**Figure 2.**
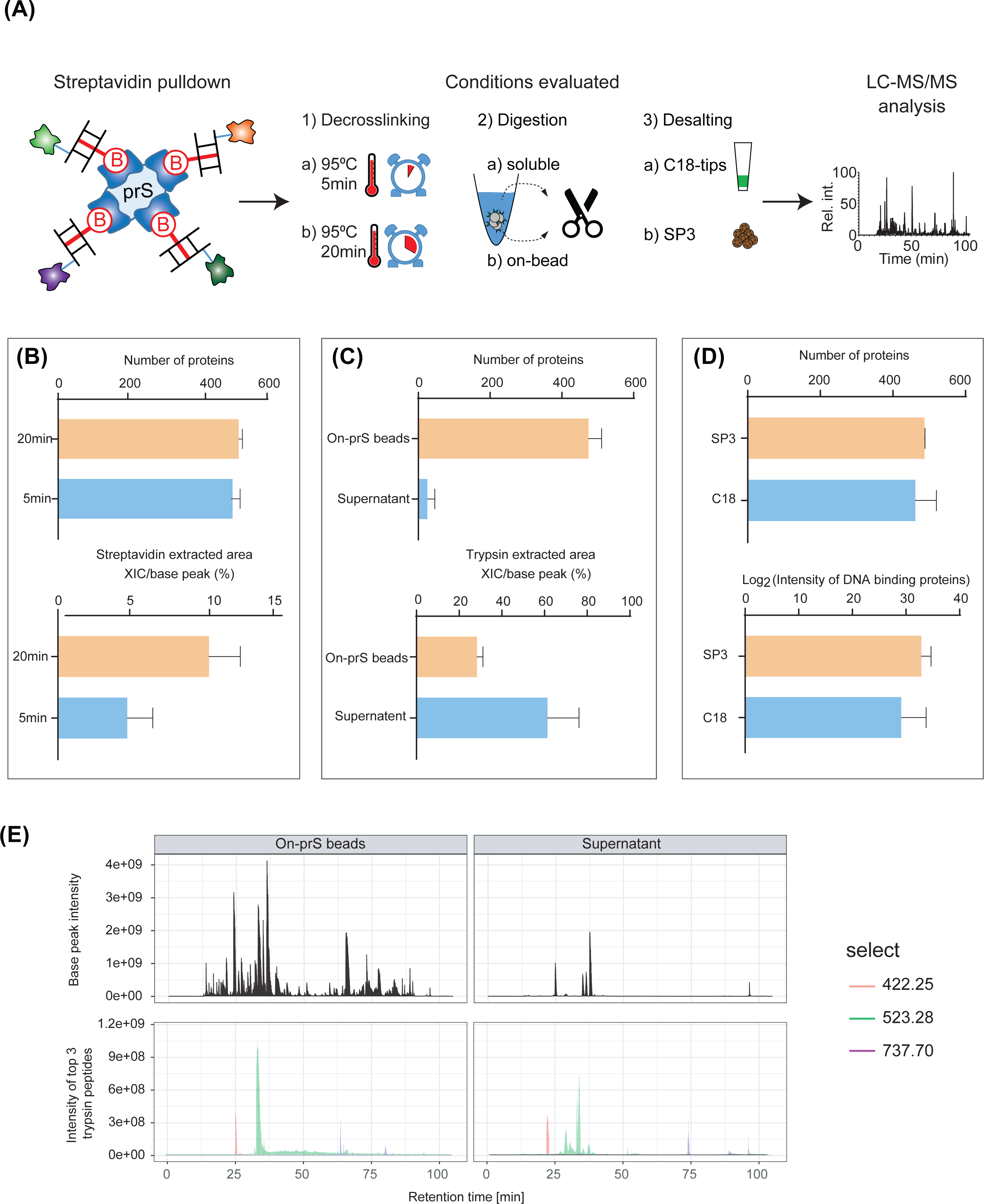
Optimization of iPOC sample processing. (A) Schematic of working strategy for the optimization of protein decrosslinking/elution, digestion, and desalting steps in the sample preparation for iPOC. (B) Number of proteins identified (top) and streptavidin percent (bottom) when heating the prS beads at 95℃ for either 20 min (orange) or 5 min (light blue). (C) Number of proteins identified (top) and trypsin percent (bottom) when performing On-prS beads (orange) or in solution (light blue) digestion. (D) Number of proteins identified (top) and Log2 transformed DNA binding proteins intensity (bottom) upon peptide clean up with either SP3 (orange) or C18 stage-tip (C18) (light blue). (E) Base peak intensity (top) and extracted ion chromatogram for the top 3 trypsin red (m/z=422.25), blue (m/z=737.70), and green (m/z=523.28) (bottom) in iPOC upon On-prS beads (left) or in solution (supernatant) (right) digestion. Relative abundance is reported on the y-axis.

### 3.3 Assessing the sensitivity of the optimized iPOC strategy *via* quantitative proteomics

To further explore the sensitivity of iPOC strategy as an *in vivo* DNA-binding protein enrichment method, we performed a competition experiment in a quantitative proteomics setting. In particular, EdU-labeled cells were subjected to iPOC in four different conditions corresponding to decreasing amounts of biotin-azide (i.e. 100%, 75%, 25%, 0%), at respective increase in competition by cy5-azide (i.e. 0%, 25%, 75%, 100%) (Fig. 3A). Western blot analyses confirmed the specificity of the click-reaction, showing the expected decrease in biotin signal upon biotin-azide competition (Fig. S3A, left). In addition, our results demonstrate a very high enrichment efficiency as measured by the low residual biotin in the flow-through of the streptavidin pull-down (Fig. S3A, right). In order to assess the sensitivity of the iPOC protocol, we designed a 10plex tandem mass tag (TMT) experiment including two biological replicates for the 4 different biotin/Cy5 proportions, plus two additional negative controls corresponding to cell extracts from cells not exposed to EdU (Fig. 3A). We compared the protein abundance obtained in the different conditions (Fig. 3B), and among the 1471 proteins identified in total, 1393 proteins (93%) showed a progressive decrease in abundance when biotin was competed with cy5-azide (Fig. 3B). This showed the high specificity and sensitivity of iPOC and suggests that these proteins may provide bona fide chromatin constituents. Indeed, these proteins were predominantly enriched in gene ontology terms associated with chromatin, (e.g. DNA-binding, chromosome organization, chromatin modification, and chromatin remodeling) (Fig. 3C). Interestingly, proteins showing unchanged intensity during Cy5 competition (Fig. 3B) are enriched in gene ontology terms associated with cytosol and RNA binding (Fig. S3B), providing a strong basis to classify them as contaminant proteins. For instance, we evaluated the intensity of different classes of DNA-binding proteins including histones (i.e. histone H1 complex), transcription factors (i.e. General transcription factor II), and chromatin assembly factors (i.e. Chromodomain helicase DNA binding protein 3), and observed the expected decreased intensity when biotin was competed with cy5-azide in iPOC (Fig. 3D). GO analysis for 94 proteins with unchanged intensity at increased biotin concentration showed that 38 proteins were located in cytosol and 25 were cytoplasmatic, including cytoskeletal proteins like actin and tubulin. (Supplementary Table1). These experiments thus emphasize the specificity of iPOC in selectively enriching for DNA-binding proteins, and show that a competition experiment is an additional way to recognize and eliminate contaminants.

**Figure 3.**
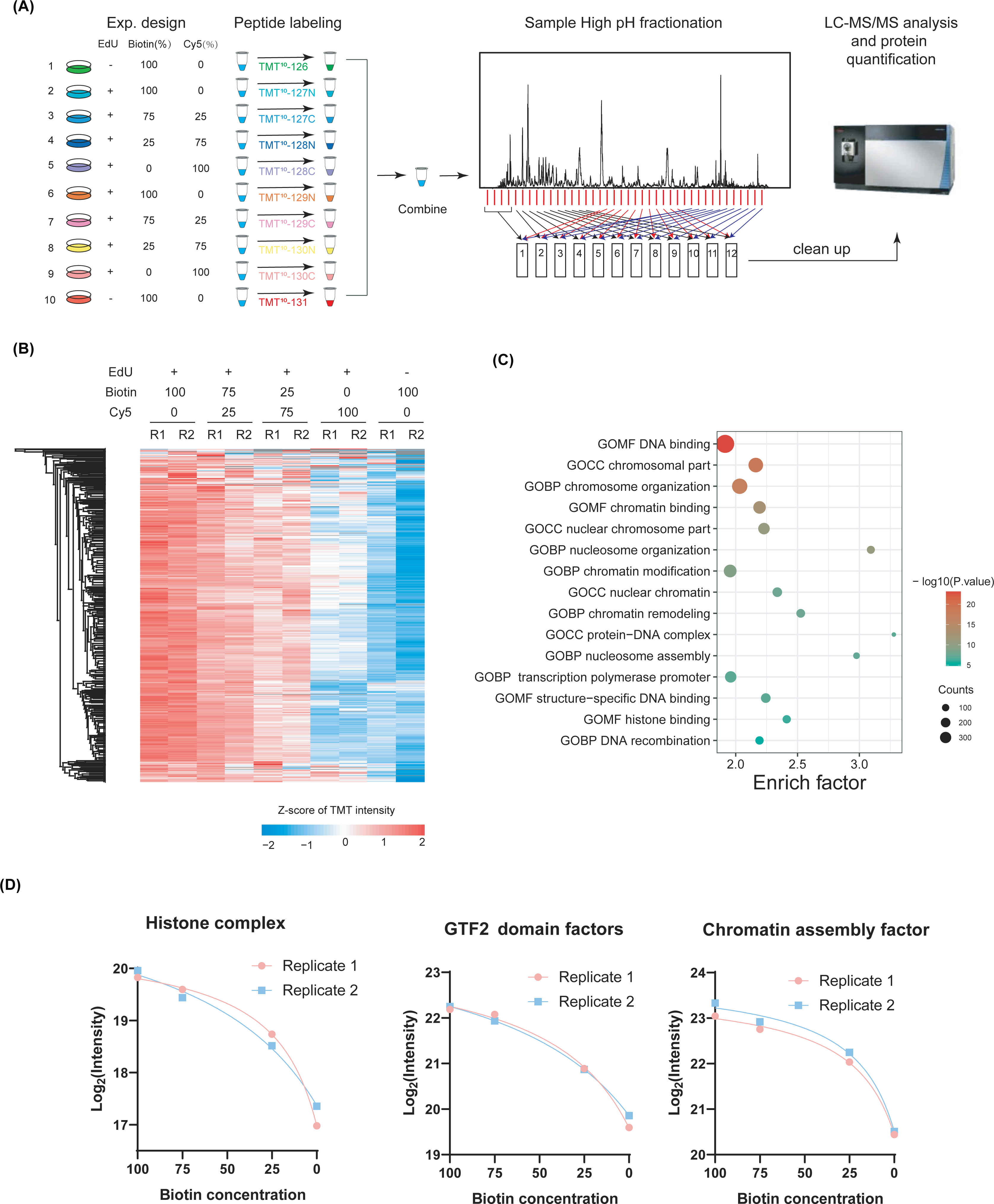
Assessment of iPOC specificity *via* TMT-mediated quantitative proteomics and Cy5-azyde competition assay. (A) Experimental design of the TMT-labeled LC-MS/MS analysis. Peptides from samples where biotin was competed at 0, 25, 75 or 100% by Cy5-azide were labeled according to TMT scheme, followed by high-pH (HpH) fractionation and LC-MS/MS quantitative analysis. (B) Heatmap representation of the intensity of proteins at different ratios between biotin- and Cy5-azide. (C) Top15 gene ontology categories associated with putative DNA-binding proteins showing decreased intensity upon biotin competition. (D) Intensity dynamics of canonical DNA-binding protein families at increasing biotin competition. Protein intensity in the two biological replicates is reported in red and blue respectively.

To benchmark the optimized iPOC protocol, we compared our result with recently developed methods for studying chromatin composition (i.e. Chromatin pelleting [32], DmChP [22], ChEP [20]), and evaluated their specificity and sensitivity in isolating chromatin proteins from cell extracts. In order to obtain a comprehensive view on the specificity of each approach, it is crucial to consider not only the total number of identified proteins, but more importantly the proportion of DNA-binding proteins identified by each strategy. Normalised to the total number of identified proteins, iPOC performed favorably compared to the other methods in capturing nuclear and DNA-binding proteins, accounting for almost 80 and 40% of all proteins identified, respectively (Fig S3C, D). This contrasts sharply with chromatin pelleting, which identified many more proteins (4000 vs 1400, Fig S3C) however retrieving a much smaller fraction of nuclear and DNA-binding proteins (Fig S3D), pointing to a substantial contamination of proteins not related to chromatin. Taken together, our results demonstrate that iPOC is a highly selective method for capturing DNA-mediated interactions and studying the composition of the DNA-bound proteome.

### 3.4 Application of iPOC to study chromatin effectors of PI3K, AKT, and mTOR signaling

We next aimed to investigate if iPOC can be used to characterize the dynamics of protein-DNA interactions downstream of a signaling cascade. Indeed, very little is yet known on how deregulation of oncogenic signaling pathways results in the association or dissociation of specific protein sets in the chromatin context, thereby regulating fundamental cellular processes ranging from cell division and motility, to transcriptional control and regulation of the cell cycle. To fill this knowledge gap, we applied iPOC to investigate how DNA-bound proteome is affected upon selective signaling inhibition. We first evaluated the efficacy of Taselisib, MK2206, and Torin 1, to verify the level of signaling activation. Upon treatment of MCF7 cells with either of these inhibitors, the phosphorylation level of PI3K, AKT, and mTOR was strongly decreased, while the total protein level stayed stable, thereby confirming efficient inhibition of the signaling pathway (Fig. 4A). It is well established that PI3K-mediated AKT activation promotes the phosphorylation of FOXO1, causing its nuclear exclusion and down-regulation of genes associated with autophagy and apoptosis [33, 34]. In line with this, upon inhibition of PI3K, AKT or mTOR, we observed decreased level of phosphorylated FOXO1 (p-FOXO1) and increased level of total FOXO1, thereby confirming the specific inhibition of these pathways (Fig. 4A). To gain a better understanding of how these signaling pathways regulate the association of proteins to DNA, we designed a SILAC-based quantitative experiment where cells labeled with intermediate or heavy channels were exposed to EdU, and the latter were additionally treated with the selective inhibitor. Light-labeled cells served as internal control (Fig. 4B). Across the three inhibitors, we identified in total 1136 proteins, of which 421 were shared among PI3Ki-, AKTi-, and mTORi-treated samples (Fig. S4A). We first benchmarked iPOC against the MS-based proteomic analysis of the chromatin input. As we previously observed [24], despite the almost 4000 proteins identified in the chromatin input (Fig. S4B), for each of the inhibitors, we identified more than 200 proteins in iPOC that were not identified in the input (Fig. S4C), thereby demonstrating the superior sensitivity of iPOC in discovering the DNA-bound proteome, as result of the selective enrichment of this class of proteins.

**Figure 4.**
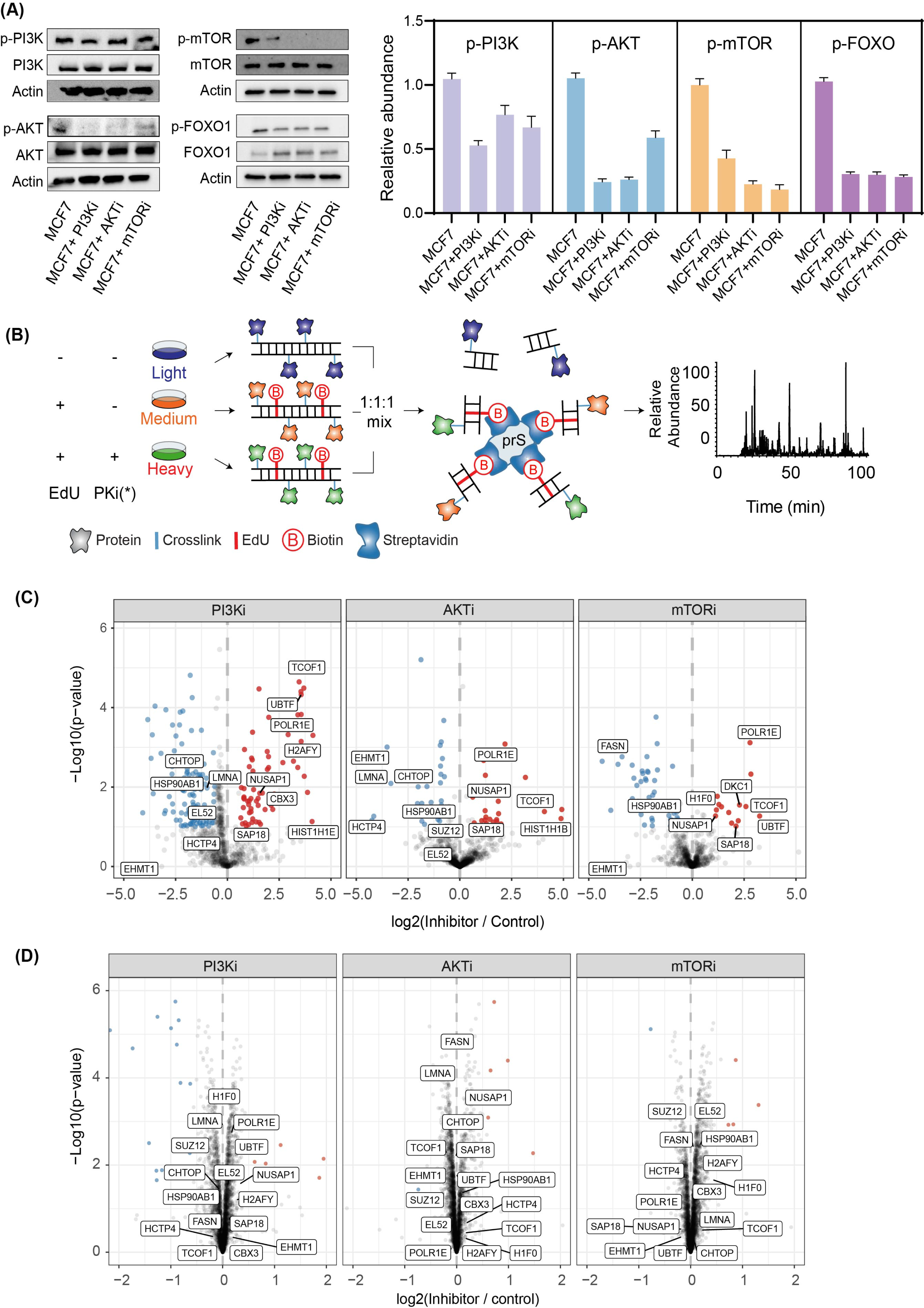
Application of iPOC to the study of PI3K, AKT, or mTOR signaling pathways (A) Western blot of phosphorylation and total level of PI3K, AKT, mTOR and FOXO upon different inhibitors. The histogram shows the corresponding quantification data. Actin was used as a loading control for these analyses. Each experiment was performed in biological triplicates and results are presented as mean ± S.E.M. (B) Schematic of iPOC experimental strategy. Light-SILAC cells served as negative control and were only treated with DMSO. Medium-SILAC cells were subjected to EdU labeling and heavy-SILAC cells were treated with both EdU and inhibitor. PKi(*) means inhibitors correspond to PI3K inhibitor Taselisib (PI3Ki), AKT inhibitor MK2206 (AKTi), or mTOR inhibitor Torin 1 (mTORi). Volcano plots showing differential expression of DNA binding proteins in (C) iPOC and (D) Input upon PI3K, AKT, or mTOR inhibition.

To identify DNA-binding proteomic features that are common or unique among the different inhibitors, we performed comparative quantitative analysis. In particular, we observed that the abundance of 161, 63 and 64 proteins was significantly deregulated upon treatment with PI3K, AKT or mTOR inhibitor, respectively (Fig. 4C, Supplementary Table2). A similar analysis performed on the chromatin input preparation only identified a limited number of deregulated proteins (Fig. 4D), thus strongly indicating that the observed patterns in iPOC are due to drug-induced changes in DNA-binding, not in protein expression. Hence, iPOC elucidates this regulatory layer that cannot be obtained from global proteome analysis.

### 3.5 Quantitative DNA-bound proteome characterization upon PI3K, AKT, or mTOR inhibition

In the signaling cascade under investigation, PI3K is known to be upstream of AKT, which in turn activates mTOR. Unsupervised hierarchical clustering of protein dynamics identified *via* iPOC shows how the DNA-bound proteome reflects this order, where inhibition of either mTOR or AKT elicit a more similar DNA-bound proteome dynamics in comparison to PI3K (Fig. S5A). This result suggests how investigation of the DNA-bound proteome *via* iPOC could potentially contribute to unravel the sequential order of proteins belonging to less-characterized signaling pathways.

In line with the hierarchical clustering, inhibition of PI3K, AKT, or mTOR results in a prevalence of proteins evicted from the DNA (Fig. S5B). This impacts a particular set of candidates: while PI3Ki treatment affects proteins involved in the spliceosome and SUMOylation, AKT inhibition deregulates the association of targets participating in the response to interleukin and regulation of hydrolase activity, and mTORi modulates proteins that regulate the cell cycle and mRNA processing (Fig. 5A-C). Interestingly, the majority of these proteins do not show a deregulation in the chromatin input (Fig. 4D), thus suggesting how the selective drug inhibition alters their DNA association without affecting their overall abundance. Although these targets differ between the treatments, they have in common that they are involved in chromosome organization and epigenetic gene regulation (Fig. 5A-C).

**Figure 5.**
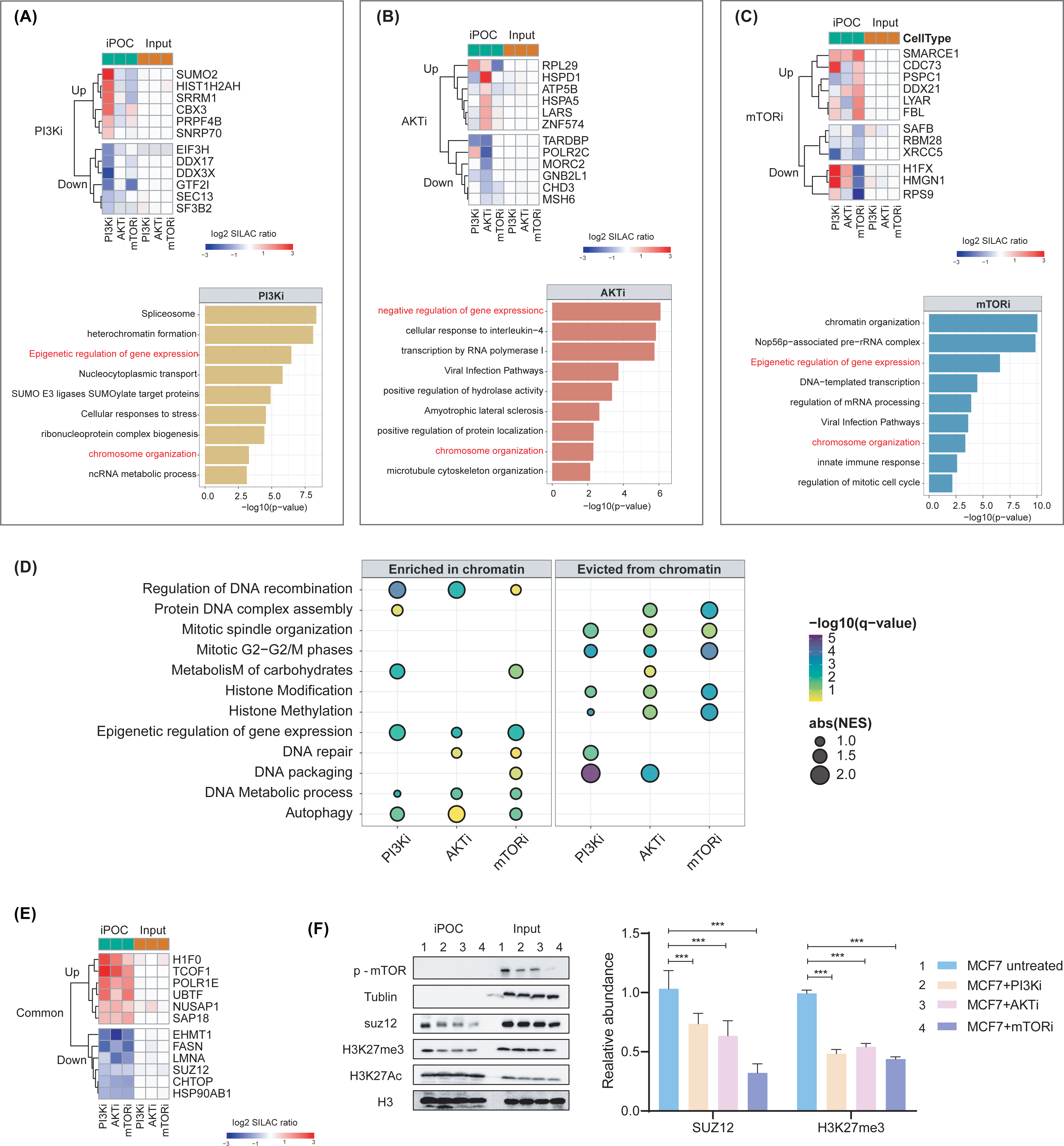
Comparative quantitative analysis of DNA-bound proteome upon PI3K, AKT or mTOR inhibitors. Heatmap and Gene ontologies terms of unique protein associated with (A) PI3K, (B) AKT or (C) mTOR inhibitors. (D) GSEA of DNA-bound proteome upon PI3K, AKT or mTOR inhibitors. (E) Heatmap of proteins commonly deregulated in iPOC and the chromatin input (Input) after treatment with PI3K, AKT, or mTOR inhibitors. (F) Western blots for p-mTOR, SUZ12, H3K27me3 in iPOC and input after MCF7 cells treated with PI3K/AKT/mTOR inhibitors. H3 and tubulin served as a loading control. The histogram shows the corresponding quantification data. Each experiment was performed in technical triplicates and results are presented as mean ± S.E.M. (*** means p < 0.01)

We next performed GSEA analysis to investigate the function of proteins showing differential DNA association induced by the different inhibitors (Fig. 5D). Among the commonly deregulated gene sets, proteins involved in DNA repair were significantly recruited at chromatin upon signaling inhibition, in line with AKT and mTOR being responsible for reducing the nuclear *foci* formation of BRCA1 and RAD51 in BRCA-proficient cells [35]. Similarly, in response to each of the inhibitors, autophagy-related proteins increased their DNA binding, while this was decreased for proteins belonging to the epigenetic regulation of gene expression (Fig. 5D). All together these results are in line with the active role of the PI3K/AKT/mTOR signaling in promoting cell survival and proliferation.

We then focused on specific proteins whose DNA association was equally modulated by the inhibition of PI3K, AKT, or mTOR (Fig. 5E, Fig. S5B). Among these, we observed the increased DNA-association of SAP18, a subunit of the SIN3-repressing complex, which facilitates histone deacetylation and transcriptional repression upon PI3K/AKT/mTOR signaling inhibition [36]. Together with the increased DNA binding of histone H1 (H1F0), this suggests a compensatory chromatin response to inhibit PI3K/AKT/mTOR signaling. In addition, we observed that three proteins involved in transcription of rDNA increased their DNA association upon either the inhibitors, namely the polymerase I subunit RPA49 (POLR1E), the treacle protein (TCOF1), and the nucleolar transcription factor 1 (UBTF) (Fig. 5D). Although decreased rDNA gene expression has been observed upon mTORi, increased stability for these proteins was reported upon PI3K/AKT/mTOR inhibition [37]. This result is in line with the stable profile observed for POLR1E, TCOF1, and UBTF in our chromatin input, while the increased DNA binding observed in iPOC suggests a relocalization of these proteins to DNA upon inhibition of the signaling pathway (Fig. 5D). Among the proteins evicted from the DNA after signaling inhibition, we identified the histone lysine methyltransferase EHMT1, and the chromatin target of protein arginine methyltrasferase 1 (CHTOP). In view of their role in gene transcription, the decreased DNA binding of EHMT1 and CHTOP suggests their active involvement in mediating the downstream transcriptional response of the PI3K/AKT/mTOR cascade[38, 39]. Another protein showing reduced DNA association upon signaling inhibition was SUZ12 (Fig. 5E), a core component of the polycomb repressive complex 2 (i.e. PRC2), responsible for catalyzing the trimethylation of K27 of histone H3[40]. Previous work has elucidated a link between inhibition of AKT or mTOR, destabilization of the PRC2 core component EZH2 at mRNA and protein level, and decrease in H3K27me3 [41]. Our results are in line with these pieces of evidence, but also suggest that inhibition of PI3K, AKT, or mTOR promotes the eviction of SUZ12 from the DNA and epigenetic regulation of chromatin *via* decrease in H3K27me3. We therefore first validated the results obtained in iPOC by evaluating the expression level and DNA-association of SUZ12 and its product H3K27me3 upon PI3K, AKT, or mTOR inhibition. Our results show that silencing of either of the targets specifically decreased the DNA-bound levels of SUZ12 and H3K27me3 (Fig. 5F), while its global expression was not significantly affected (Fig. S5C). The only exception was the decreased global level of H3K27me3 upon mTOR inhibition, in line with the role of mTORC2 in preventing the production of S-adenosylmethionine, a common donor of methyl groups [41].

All together our results show for the first time the chromatin dynamics upon inhibition of PI3K/AKT/mTOR pathway, and propose a novel mechanism where SUZ12 bridges the epigenetic regulation in response to signaling cascade perturbation. In addition, our data demonstrate how iPOC is a sensitive approach to identify novel DNA-bound dynamics, thereby increasing our knowledge on chromatin regulation.

## Discussion

In recent years, several methods have been established to capture, identify, and quantify DNA-binding proteins in order to investigate chromatin dynamics in time and space [19]. Among these, incorporation of EdU has been efficiently exploited to characterize the chromatin composition at defined genomic regions[21, 22, 42]. Our laboratory has recently widened this view point and developed iPOC, a strategy to investigate the global chromatin dynamic changes elicited during the DNA damage repair [24]. The iPOC approach combines SILAC labeling for the robust protein quantification, with capture of biotinylated DNA-protein complexes on protease-resistant streptavidin (prS) beads [25], thus highly increasing the sensitivity of the approach. In order to boost the enrichment efficiency and the signal to noise, here we have optimized a number of key steps in the iPOC sample preparation procedure with regard to EdU concentration, prS beads-to-input ratio, time for decrosslinking, protein digestion, and peptide desalting strategies, thereby establishing the optimal working conditions to study the DNA-bound proteome.

So far, the regulation of signaling cascades is often investigated at the level of protein phosphorylation, either using phospho-specific antibodies in either ELISA or western blots to confirm pathway activation [43], or by phospho-proteomics to investigate kinase-dependent signaling networks in a more unbiased and integrative fashion [44]. Despite their power and wide-spread use, these approaches primarily capture upstream events occurring in the cytosol upon pathway activation, while they are not designed to describe downstream consequences in chromatin, representing an important regulatory layer that mediates signaling output (i.e. gene expression). Various methods have been designed to investigate chromatin composition[21–23], however none of them have been applied in the context of signaling cascades. In addition, strategies based on proximity biotinylation have been explored to identify proteins that interact or colocalize with a selected target of interest in chromatin [45], however they require the engineering of a pre-defined protein of interest fused to a biotin donor, such as BirA or APEX [46, 47]. In contrast, iPOC does not require cellular engineering nor *a priori* knowledge on a specific target protein, and uses DNA as the focal point to study DNA-bound interactors (chromatome). We previously showed the instrumental role of iPOC to investigate chromatome changes during DNA damage repair [24], and we here refined and applied our methodology to study the chromatome dynamics upon deregulation of mitogenic signaling. To the best of our knowledge, this is the first time that a chromatin-directed strategy has been used to globally investigate alterations in the DNA-bound proteome elicited by a signaling pathway, and more generally opens the way to study chromatin as a regulatory layer between upstream signaling and downstream cellular response.

With iPOC, we studied the dynamics of the DNA-bound proteome upon treatment with PI3K, AKT, and mTOR inhibitors, and in agreement with previous studies we observed that the DNA-bound state of proteins involved in autophagy and chromatin organization is globally modulated. In addition, our comparative analysis revealed that a subset of proteins involved in epigenetic regulation of gene expression was commonly deregulated upon inhibition of either the targets. Among these, the increase in the DNA-bound fraction of SAP18 corroborates previous observations on the antagonistic mechanism between the SIN3-repressing complex and PI3K[48]; moreover our results indicate that AKT and mTOR might also have a similar effect on SAP18. Furthermore, the reduced DNA binding of CHTOP (chromatin recruiter of PRMT1) and the methyltrasferase EHMT1 is in line with the decreased methylation of specific chromatin targets associated with decrease in cell proliferation and DNA replication, and increased cell apoptosis, respectively[48, 49].

In addition, in iPOC we observed that the association of SUZ12 with the DNA was reduced upon inhibition of PI3K/AKT/mTOR signaling pathway. SUZ12 catalyzes the trimethylation of lysine 27 on histone H3 (H3K27me3), a known marker of poised or silenced promoters, that strongly influences tumor pregression and malignancy [40]. Recent studies have shown that mTOR signaling promotes proliferation of glioblastoma cells through the induction of H3K27me3, and EZH2 has been proposed as mediator of such epigenetic rewiring in multiple myeloma [40, 41]. Our results provide a novel view, and suggests that PI3K/AKT/mTOR signaling might regulate the DNA binding status of SUZ12, thereby modulating the epigenetic cell state. A recent report on the responsiveness of JAZF1-SUZ12 fusion sarcomas to mTOR inhibition [50] further supports our hypothesis, thereby suggesting SUZ12 as an effective drug target in cancers with hyperactive PI3K/AKT/mTOR signaling. Thus, iPOC provides a novel view on how this cascade could modulate the cell epigenetic status, and suggests a mechanism through which oncogenic pathway deregulation is translated into a cancer-specific chromatome and epigenetic status. Further functional studies will be required to corroborate these hypotheses.

Although belonging to the same signaling cascade, application of iPOC to the study of the PI3K/AKT/mTOR pathway shows that the selective inhibition of either the targets impinges on the DNA-bound state of a rather specific set of downstream effectors. These observations enhance our global understanding of how these three components of the PI3K/AKT/mTOR pathway trigger the stimulus to the DNA. Moreover, further characterization of DNA-binding proteins specifically regulated by either the proteins may also identify novel therapeutic targets to be potentially exploited in drug resistance often arising in mono-therapies.

In conclusion, iPOC is an unbiased and sensitive approach to identify the dynamic chromatin association even for transiently interacting DNA-binding proteins, and therefore we anticipate that iPOC will be a useful tool to increase mechanistic understanding of how signaling pathways impinge on the DNA.

## Data availability

The mass spectrometry proteomics data and the search data by MaxQuant (version 1.6.1.0) have been deposited to the ProteomeXchange Consortium via the PRIDE partner repository [51] with the dataset identifier PXD045235. (Reviewer account details: Username, Password: pBxhULwn).

## Acknowledgements

We thank the China Scholarship Council awarded a scholarship to Huiyu Wang, so that she can pursue study in Germany as a Visiting PhD Student. We also thank Dr. Hua xiao from the School of Life Sciences and Biotechnology at Shanghai Jiao Tong University for supporting Huiyu Wang to study in Germany.

## Reference

1. Aranda S. and Di Croce L., Isolation of Chromatin Proteins by Genome Capture. Methods Mol Biol, 2023. 2655:91–99.

2. Bonev B. and Cavalli G., Organization and function of the 3D genome. Nat Rev Genet, 2016. 17(11):661–678.

3. Espejo I., Di Croce L. and Aranda S., The changing chromatome as a driver of disease: A panoramic view from different methodologies. Bioessays, 2020. 42(12):e2000203.

4. Clapier C.R. and Cairns B.R., The biology of chromatin remodeling complexes. Annu Rev Biochem, 2009. 78:273–304.

5. Mirabella A.C., Foster B.M. and Bartke T., Chromatin deregulation in disease. Chromosoma, 2016. 125(1):75–93.

6. Vaughan R.M., Kupai A. and Rothbart S.B., Chromatin Regulation through Ubiquitin and Ubiquitin-like Histone Modifications. Trends Biochem Sci, 2021. 46(4):258–269.

7. Bailey M.H., Tokheim C., Porta-Pardo E., et al., Comprehensive Characterization of Cancer Driver Genes and Mutations. Cell, 2018. 174(4):1034–1035.

8. Kandoth C., McLellan M.D., Vandin F., et al., Mutational landscape and significance across 12 major cancer types. Nature, 2013. 502(7471):333-+.

9. Martin C.J. and Moorehead R.A., Polycomb repressor complex 2 function in breast cancer. Int J Oncol, 2020. 57(5):1085–1094.

10. Pfister S.X. and Ashworth A., Marked for death: targeting epigenetic changes in cancer. Nat Rev Drug Discov, 2017. 16(4):241–263.

11. Glaviano A., Foo A.S.C., Lam H.Y., et al., PI3K/AKT/mTOR signaling transduction pathway and targeted therapies in cancer. Mol Cancer, 2023. 22(1).

12. Li Q., Li Z., Luo T., et al., Targeting the PI3K/AKT/mTOR and RAF/MEK/ERK pathways for cancer therapy. Mol Biomed, 2022. 3(1):47.

13. Tewari D., Patni P., Bishayee A., et al., Natural products targeting the PI3K-Akt-mTOR signaling pathway in cancer: A novel therapeutic strategy. Semin Cancer Biol, 2022. 80:1–17.

14. Zhao J., Zhai B., Gygi S.P., et al., mTOR inhibition activates overall protein degradation by the ubiquitin proteasome system as well as by autophagy. Proc Natl Acad Sci U S A, 2015. 112(52):15790–15797.

15. Jhaveri K., Chang M.T., Juric D., et al., Phase I Basket Study of Taselisib, an Isoform-Selective PI3K Inhibitor, in Patients with PIK3CA-Mutant Cancers. Clin Cancer Res, 2021. 27(2):447–459.

16. Sangai T., Akcakanat A., Chen H.Q., et al., Biomarkers of Response to Akt Inhibitor MK-2206 in Breast Cancer. Clin Cancer Res, 2012. 18(20):5816–5828.

17. Bin Emran T., Shahriar A., Mahmud A.R., et al., Multidrug Resistance in Cancer: Understanding Molecular Mechanisms, Immunoprevention and Therapeutic Approaches. Front Oncol, 2022. 12.

18. Ou H.D., Phan S., Deerinck T.J., et al., ChromEMT: Visualizing 3D chromatin structure and compaction in interphase and mitotic cells. Science, 2017. 357(6349).

19. Sigismondo G., Papageorgiou D.N. and Krijgsveld J., Cracking chromatin with proteomics: From chromatome to histone modifications. Proteomics, 2022. 22(15-16).

20. Kustatscher G., Wills K.L.H., Furlan C., et al., Chromatin enrichment for proteomics. Nat Protoc, 2014. 9(9):2090–2099.

21. Sirbu B.M., Couch F.B. and Cortez D., Monitoring the spatiotemporal dynamics of proteins at replication forks and in assembled chromatin using isolation of proteins on nascent DNA. Nat Protoc, 2012. 7(3):594–605.

22. Aranda S., Alcaine-Colet A., Blanco E., et al., Chromatin capture links the metabolic enzyme AHCY to stem cell proliferation. Sci Adv, 2019. 5(3).

23. Aranda S., Borràs E., Sabidó E., et al., Chromatin-Bound Proteome Profiling by Genome Capture. Star Protocols, 2020. 1(1).

24. Sigismondo G., Arseni L., Palacio-Escat N., et al., Multi-layered chromatin proteomics identifies cell vulnerabilities in DNA repair. Nucleic Acids Res, 2023. 51(2):687–711.

25. Rafiee M.R., Sigismondo G., Kalxdorf M., et al., Protease-resistant streptavidin for interaction proteomics. Mol Syst Biol, 2020. 16(5).

26. Hughes C.S., Moggridge S., Müller T., et al., Single-pot, solid-phase-enhanced sample preparation for proteomics experiments. Nat Protoc, 2019. 14(1):68–85.

27. Rappsilber J., Ishihama Y. and Mann M., Stop and go extraction tips for matrix-assisted laser desorption/ionization, nanoelectrospray, and LC/MS sample pretreatment in proteomics. Anal Chem, 2003. 75(3):663–670.

28. Cox J. and Mann M., MaxQuant enables high peptide identification rates, individualized p.p.b.-range mass accuracies and proteome-wide protein quantification. Nat Biotechnol, 2008. 26(12):1367–1372.

29. Tyanova S., Temu T. and Cox J., The MaxQuant computational platform for mass spectrometry-based shotgun proteomics. Nature Protocols, 2016. 11(12):2301–2319.

30. Zhou Y.Y., Zhou B., Pache L., et al., Metascape provides a biologist-oriented resource for the analysis of systems-level datasets. Nat Commun, 2019. 10.

31. Schneider C.A., Rasband W.S. and Eliceiri K.W., NIH Image to ImageJ: 25 years of image analysis. Nat Methods, 2012. 9(7):671–675.

32. van Mierlo G., Dirks R.A.M., De Clerck L., et al., Integrative Proteomic Profiling Reveals PRC2-Dependent Epigenetic Crosstalk Maintains Ground-State Pluripotency. Cell Stem Cell, 2019. 24(1):123-+.

33. Fabre S., Lang V., Harriague J., et al., Stable activation of phosphatidylinositol 3-kinase in the T cell immunological synapse stimulates Akt signaling to FoxO1 nuclear exclusion and cell growth control. J Immunol, 2005. 174(7):4161–4171.

34. Huang H., Regan K.M., Lou Z., et al., CDK2-dependent phosphorylation of FOXO1 as an apoptotic response to DNA damage. Science, 2006. 314(5797):294–297.

35. Plo I., Laulier C., Gauthier L., et al., AKT1 Inhibits Homologous Recombination by Inducing Cytoplasmic Retention of BRCA1 and RAD51. Cancer Res, 2008. 68(22):9404–9412.

36. Hassig C.A., Fleischer T.C., Billin A.N., et al., Histone deacetylase activity is required for full transcriptional repression by mSin3A. Cell, 1997. 89(3):341–347.

37. Mayer C., Zhao J., Yuan X., et al., mTOR-dependent activation of the transcription factor TIF-IA links rRNA synthesis to nutrient availability. Genes Dev, 2004. 18(4):423–434.

38. Izumikawa K., Ishikawa H., Simpson R.J., et al., Modulating the expression of Chtop, a versatile regulator of gene-specific transcription and mRNA export. RNA Biol, 2018. 15(7):849–855.

39. Watson Z.L., Yamamoto T.M., McMellen A., et al., Histone methyltransferases EHMT1 and EHMT2 (GLP/G9A) maintain PARP inhibitor resistance in high-grade serous ovarian carcinoma. Clin Epigenetics, 2019. 11(1):165.

40. Rizk M., Rizq O., Oshima M., et al., Akt inhibition synergizes with polycomb repressive complex 2 inhibition in the treatment of multiple myeloma. Cancer Sci, 2019. 110(12):3695–3707.

41. Harachi M., Masui K., Honda H., et al., Dual Regulation of Histone Methylation by mTOR Complexes Controls Glioblastoma Tumor Cell Growth via EZH2 and SAM. Mol Cancer Res, 2020. 18(8):1142–1152.

42. Aranda S., Rutishauser D. and Ernfors P., Identification of a large protein network involved in epigenetic transmission in replicating DNA of embryonic stem cells. Nucleic Acids Res, 2014. 42(11):6972–6986.

43. Xu W., Wang Z.Q., Zhang Z., et al., PIK3CB promotes oesophageal cancer proliferation through the PI3K/AKT/mTOR signalling axis. Cell Biol Int, 2022. 46(9):1399–1408.

44. Franciosa G., Locard-Paulet M., Jensen L.J., et al., Recent advances in kinase signaling network profiling by mass spectrometry. Curr Opin Chem Biol, 2023. 73.

45. Ummethum H. and Hamperl S., Proximity Labeling Techniques to Study Chromatin. Front Genet, 2020. 11.

46. Roux K.J., Kim D.I., Raida M., et al., A promiscuous biotin ligase fusion protein identifies proximal and interacting proteins in mammalian cells. J Cell Biol, 2012. 196(6):801–810.

47. Rhee H.W., Zou P., Udeshi N.D., et al., Proteomic Mapping of Mitochondria in Living Cells via Spatially Restricted Enzymatic Tagging. Science, 2013. 339(6125):1328–1331.

48. Varier K.M., Dan G., Liu W.L., et al., Stilbene B10 induces apoptosis and tumor suppression in lymphoid Raji cells by BTK-mediated regulation of the KRAS/HDAC1/EP300/PEBP1 axis. Biomed Pharmacother, 2022. 156.

49. Giuliani V., Miller M.A., Liu C.Y., et al., PRMT1-dependent regulation of RNA metabolism and DNA damage response sustains pancreatic ductal adenocarcinoma. Nat Commun, 2021. 12(1).

50. Chiang S., Vasudevaraja V., Serrano J., et al., TSC2-mutant uterine sarcomas with fusions demonstrate hybrid features of endometrial stromal sarcoma and PEComa and are responsive to mTOR inhibition. Modern Pathol, 2022. 35(1):117–127.

51. Perez-Riverol Y., Csordas A., Bai J.W., et al., The PRIDE database and related tools and resources in 2019: improving support for quantification data. Nucleic Acids Res, 2019. 47(D1):D442–D450.

