## Supplementary info for "Isolation of proteins on chromatin (iPOC) reveals signaling pathway-dependent alterations in the DNA-bound proteome"

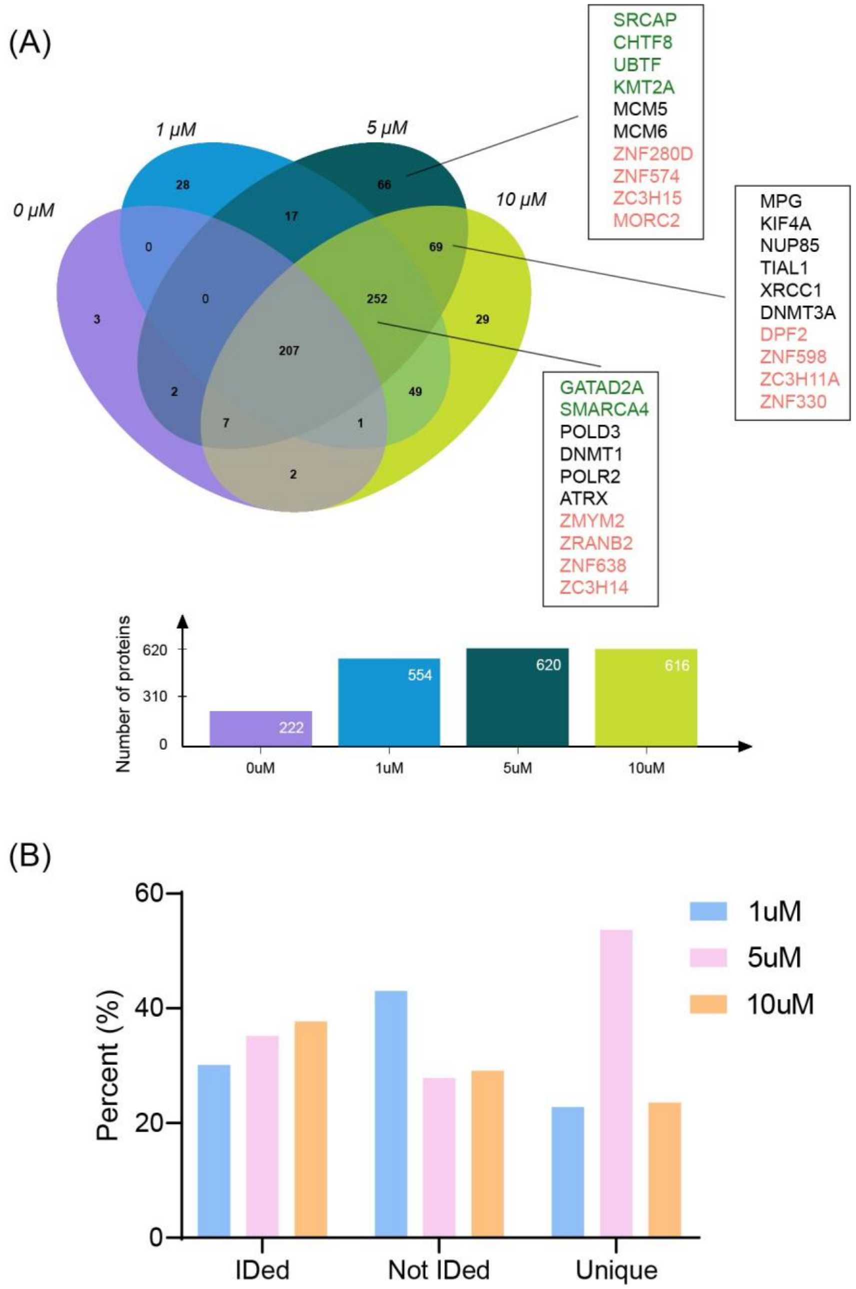


**Figure S1.** (A) Venn diagram of proteins enriched in iPOC performed with 0, 1, 5, 10μM EdU for 18h. Pink indicates proteins belonging to zinc finger protein family, green indicates transcription factors, while names in black are additional DNA-binding proteins. Histogram shows the total amount of identified proteins. (B) Bar chart represents the percent of proteins identified (IDed), not identified (Not IDed), and unique proteins in iPOC performed with 1, 5, and 10 μM EdU labeling.

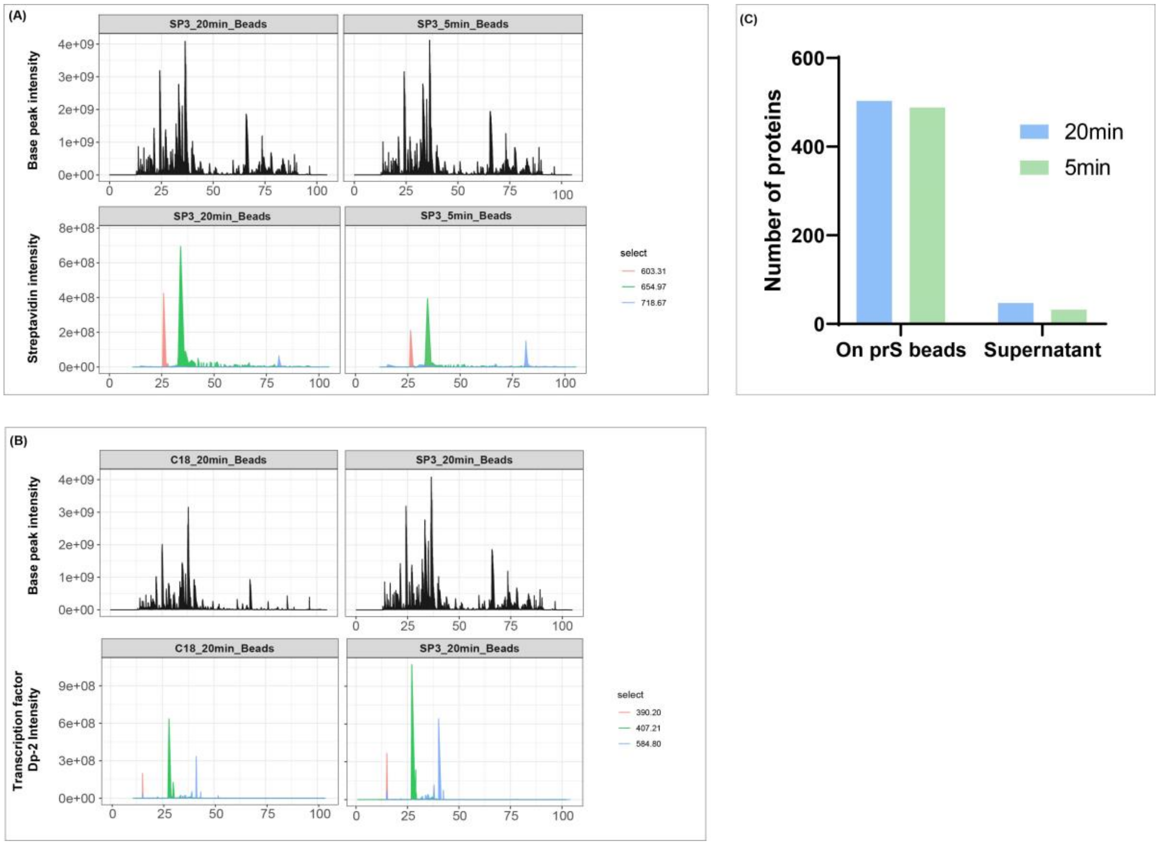


**Figure S2.** Evaluation of contaminants in chromatographic profile of iPOC samples. (A) Base peak chromatogram in iPOC experiments using 20min (left) or 5 min (right) for decrosslinking. Relative abundance is reported on the y-axis. Up is the total ion intensity, bottom is the extracted ion chromatography of the top 3 streptavidin peptides red (m/z=422.25), blue (m/z=737.70), and green (m/z=523.28). (B) Base peak chromatogram of iPOC experiment upon C18 stage-tips (C18, left) or SP3 peptide clean up. Relative abundance is reported on the y-axis. Up is the total ion intensity and down is the extracted ion chromatography of the top 3 transcription factor Dp2 peptides red (m/z=422.25), blue (m/z=737.70), and green (m/z=523.28). (C) Number of proteins identified upon digestion On prS beads or in solution (Supernatant) upon decrosslinking for 20min (light blue) or 5min (green).

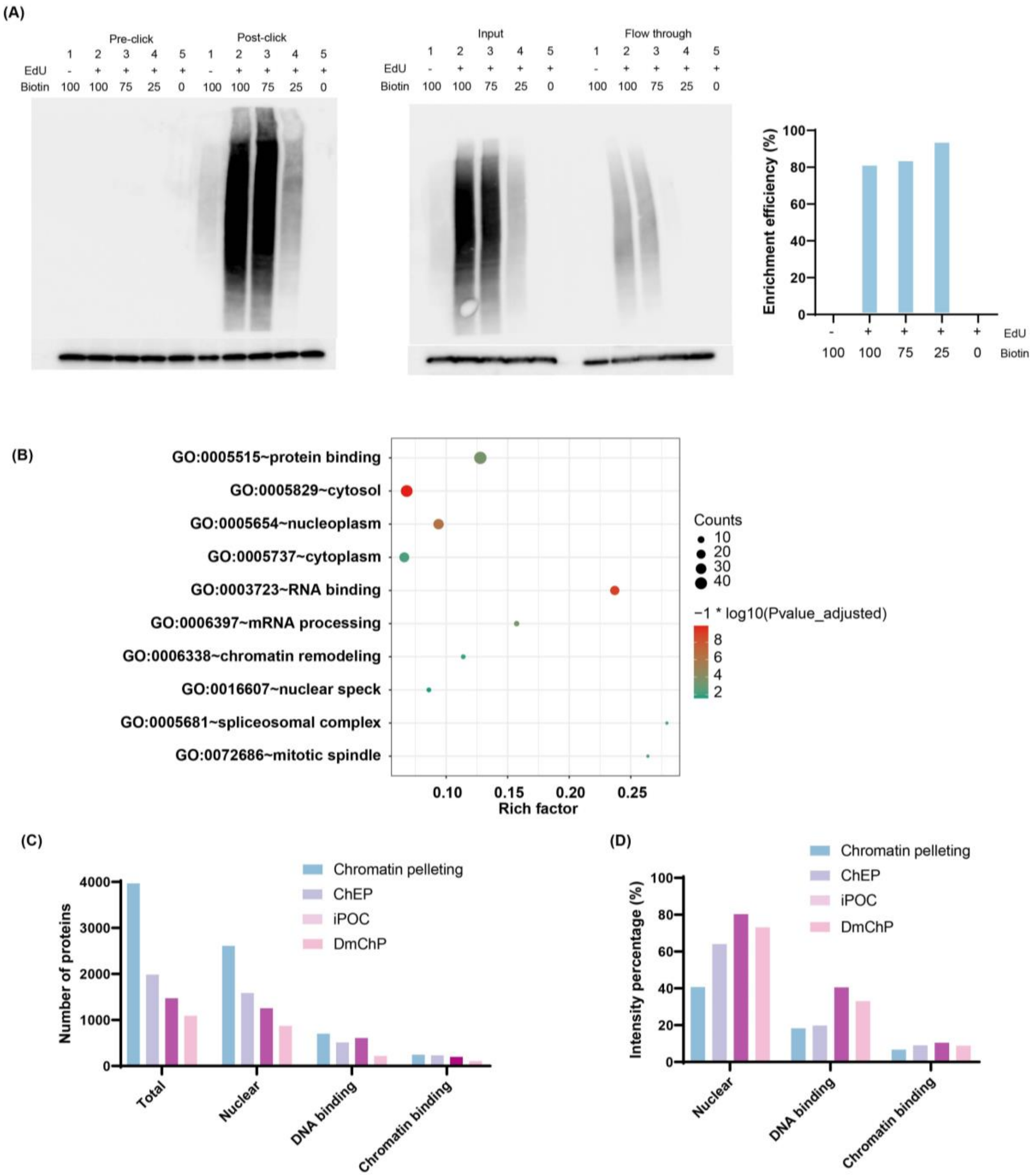


**Figure S3.** (A) Western blot of biotin (top) and histone H2A (bottom) before and after biotin azide click reaction (left) or before and after prS-mediated enrichment (center) in iPOC performed at different biotin concentrations. Bar plot (right) shows the calculated enrichment efficiency. (B) Gene ontology categories associated with proteins showing the same intensity upon biotin competition. (C) Total number of proteins and (D) percentage of total intensity for nuclear, DNA- and chromatin-binding proteins from different enrichment strategies to study DNA-binding proteins.

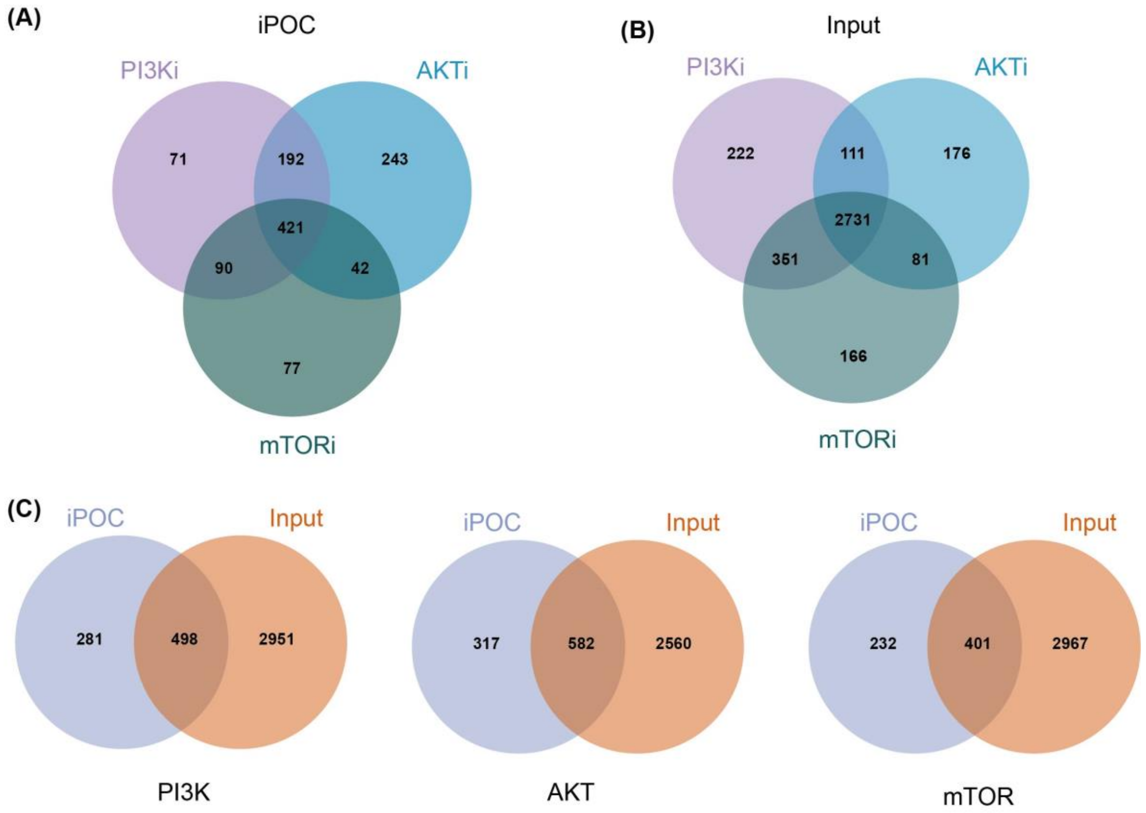


**Figure S4.** (A) Venn diagram for iPOC-derived DNA-binding proteomes. Numbers indicate the proteins identified in each of the three inhibitors. (B) Venn diagram for chromatin input proteins. Numbers indicate the proteins identified in each of the three inhibitors. (C) Venn diagram for proteins from iPOC and chromatin input (Input) for PI3K, AKT and mTOR inhibition.

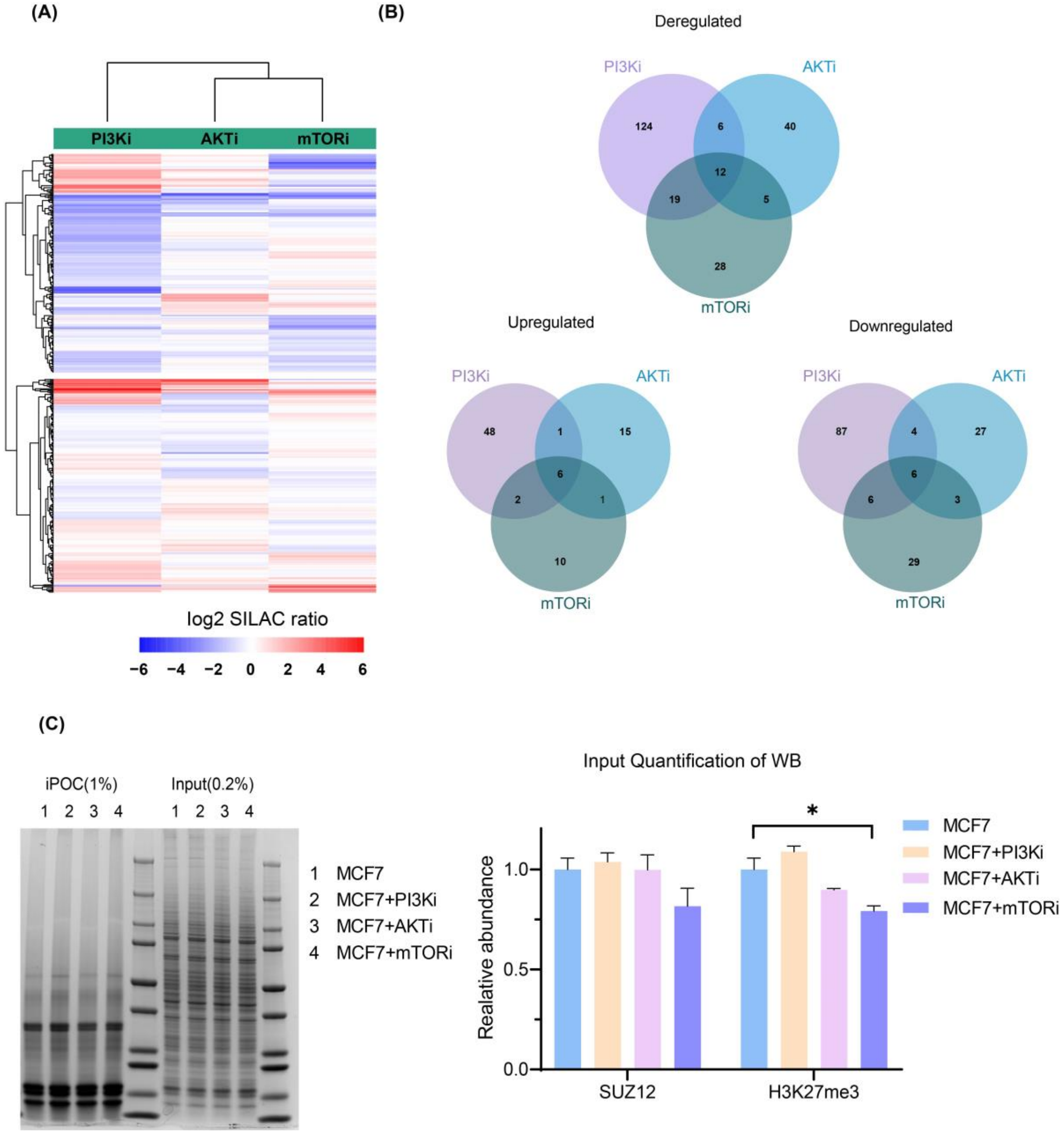


**Figure S5.** (A) Unsupervised hierarchical clustering of proteins identified in iPOC performed upon PI3K, AKT, or mTOR inhibition. (B) Venn diagram for all deregulated, upregulated and downregulated iPOC-derived DNA-binding proteomes identified upon PI3K, AKT, or mTOR inhibition. (C) Coomassie stained iPOC and chromatin input (Input) visualization for normalization (left) and western blot quantification of SUZ12 and H3K27me3 in the chromatin input (right) in cells untreated or upon selective inhibition of PI3K, AKT, or mTOR. Each experiment was performed in biological triplicates and results are presented as mean ± S.E.M. * corresponds to p-value <0.05.
